# Automated delineation of non-small cell lung cancer

**DOI:** 10.1101/304949

**Authors:** Maliazurina Saad

## Abstract

**Purpose:** The tendencies of non-small cell lung cancers (NSCLC) to be large-sized, irregularly shaped, and to grow against the surrounding structures can cause even expert clinicians to experience difficulty with accurate segmentation.

**Methods:** An automated delineation tool based on spatial analysis was developed and studied on 25 sets of CT scans of primary NSCLCs with diverse radiological characteristics (sizes, shapes, contouring, localization, and microenvironment). Manual and automated gross delineations of the gross tumor were compared using a specific metric built based on spatially overlapping pixels.

**Results:** The proposed algorithm exhibited robustness in terms of the tumor size (5.32–18.24 mm), shape (spherical or non-spherical), contouring (lobulated, spiculated, or cavitated), localization (solitary, pleural, mediastinal, endobronchial, or tagging), and microenvironment (left or right lobe), with sensitivity, specificity, and accuracy rates of 80–98%, 85–99%, and 84–99%, respectively.

**Conclusions:** Small discrepancies were observed between the manual and automated delineations. These might have risen from the variability in the practitioners’ definitions of ROIs or from imaging artifacts that reduced the tissue resolution.

## 1. Introduction

Suspicious region delineation refers to a localization phase wherein a clinician defines the exact position and border of region of interest (ROI), which may encompass a cancerous or non-cancerous pathological object requiring further analysis [1–2]. The study focuses on the initial quantitative steps in the process of clinical decision making, suspicious region delineation, for patients with non-small-cell lung cancer (NSCLC). Remarkably, this step is performed before any other analysis, leading to quantitatively deduced decisions. NSCLCs are often large-sized with an irregular shape and invasion of the surrounding structures in comparison with lung nodules. These features present challenges and cause the failure of automatic segmentation and detection algorithms developed for small lung nodules [3–7], particularly for the following reasons: (1) NSCLCs are often attached to the pleural or mediastinal wall and are easily excluded during thoracic segmentation. (2) Existing algorithms always assume a small, solid, and round lung nodule, in contrast to the characteristic features of NSCLC. (3) It may be impossible to distinguish the NSCLC from the surrounding structures with similar intensities on radiological images.

To address these limitations, expert clinicians generally delineate the NSCLCs manually with little, if any, machine assistance. However, this process is time-consuming, labor-intensive, and prone to interobserver variability. Although several studies have evaluated the agreement between automated machine-assisted and manual contouring by humans [8–10], the former is frequently unable to match the accuracy of the latter, which remains universally acknowledged as a gold standard. Furthermore, little to no information is available regarding the robustness of the proposed tools in dealing with the mentioned issues. Therefore, this study aims to propose a delineation tool with sufficient robustness to address all possible radiological conditions and reduce the accuracy gap between the automated and manual delineations.

## 2. Materials and Methods

### 2.1 Subjects

25 sets of CT scans curated from the NSCLC-Radiomics dataset in the Cancer Imaging Archive [11], which were made available for download by the Dana–Farber Cancer Institute in accordance with the Dutch law and Institutional Review Board approval [12]. The patients were selected based on the inclusion criteria presented in Table 1, and the patients with multiple tumors were excluded. Manual delineation of the 3D gross tumor volume and clinical outcome data are available for the pretreatment CT scans of all patients. The data acquisition protocols varied slightly among patients, depending on their body size. The reconstructed pixel resolution was 0.977 × 0.977 mm, and all images were reconstructed using 512 × 512 pixel matrices. In all examinations, the slice thickness was 3.0 mm.

**Table 1.** Radiological characteristics of tumors. Their representation is shown in Fig. 1

| Patient ID | Size<br>(mm) | Contouring |  |  | Shape |  | Microenvironment<br>(lobe) |  |
| --- | --- | --- | --- | --- | --- | --- | --- | --- |
|  |  | Lobulated | Cavitated | Spiculated | Spherical | Non-Spherical | Left | Right |
| Solitary Mass |  |  |  |  |  |  |  |  |
| Patient 015 | 5.32 |  |  | √ | √ |  |  | √ |
| Patient 024 | 9.53 | √ |  |  | √ |  |  | √ |
| Patient 051 | 14.82 | √ |  |  | √ |  |  | √ |
| Patient 078 | 8.42 | √ |  |  | √ |  |  | √ |
| Patient 252 | 6.57 |  |  | √ | √ |  | √ |  |
| Mediastinal Mass |  |  |  |  |  |  |  |  |
| Patient 026 | 14.52 |  |  | √ |  | √ |  | √ |
| Patient 092 | 15.4 | √ |  |  |  | √ |  | √ |
| Patient 109 | 17.13 |  | √ |  |  | √ |  | √ |
| Patient 123 | 14.32 |  |  | √ |  | √ |  | √ |
| Patient 220 | 12.59 | √ |  |  | √ |  | √ |  |
| Pleural Mass |  |  |  |  |  |  |  |  |
| Patient 070 | 8.68 |  | √ |  |  | √ | √ |  |
| Patient 211 | 10.09 |  |  | √ | √ |  | √ |  |
| Patient 236 | 9.82 | √ |  |  | √ |  |  | √ |
| Patient 243 | 8.73 |  |  | √ |  | √ |  | √ |
| Patient 318 | 7.96 |  |  | √ |  | √ |  | √ |
| Endobronchial Mass |  |  |  |  |  |  |  |  |
| Patient 037 | 6.32 |  |  | √ |  | √ |  | √ |
| Patient 041 | 13.22 | √ |  |  | √ |  |  | √ |
| Patient 047 | 8.59 |  |  | √ |  | √ |  | √ |
| Patient 064 | 9.08 | √ |  |  | √ |  |  | √ |
| Patient 350 | 7.35 | √ |  |  |  | √ | √ |  |
| Tagging Mass |  |  |  |  |  |  |  |  |
| Patient 008 | 12.4 |  |  | √ | √ |  |  | √ |
| Patient 035 | 18.25 |  | √ |  |  | √ | √ |  |
| Patient 054 | 13.55 |  | √ |  |  | √ |  | √ |
| Patient 224 | 16.14 |  | √ |  |  | √ |  | √ |
| Patient 297 | 12.94 |  |  | √ |  | √ |  | √ |

**Fig. 1.**
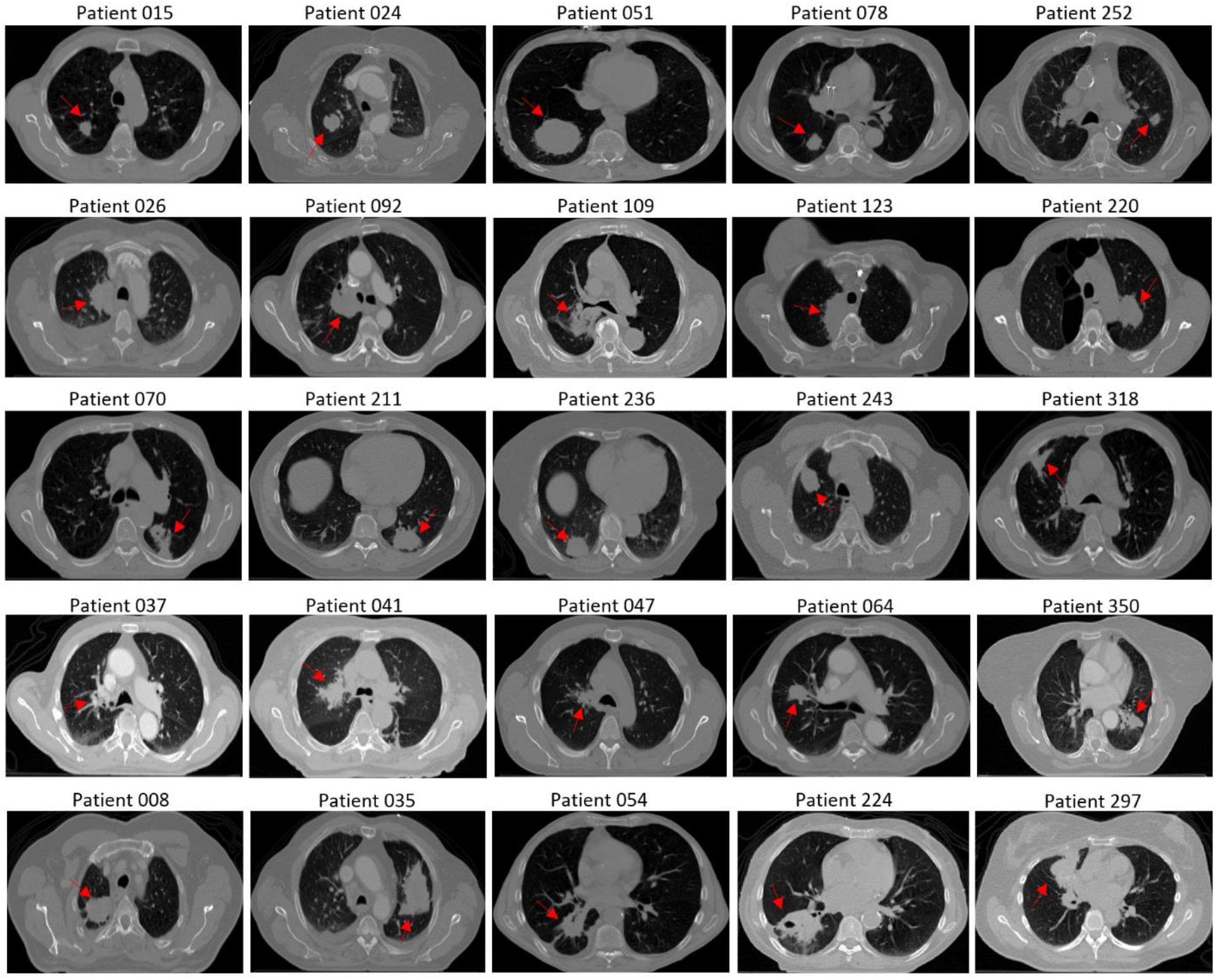
The representation of tumor as described in Table 1. Patients with solitary, mediastinal, pleural, endobronchial and tagging tumor represented by the first, second, third, fourth and fifth row respectively.

### 2.2 Framework description

An image segmentation based on a series of spatial-related techniques as shown in Fig. 2 is proposed. First, a prior processing using two-fold thresholding was first applied. Two binary masks were produced at this stage. The optimal thresholding discussed by Nihad et al. [13] were slightly modified as follows: when the threshold value T_opt_ converged, the value was applied twice to an image I as lower-bound (M_LB_) and upper-bound masks (M_UB_), as depicted in Eq.1 and Eq.2, respectively.

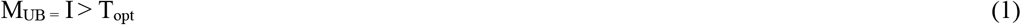

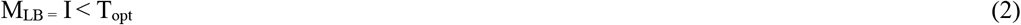

**Fig. 2.**
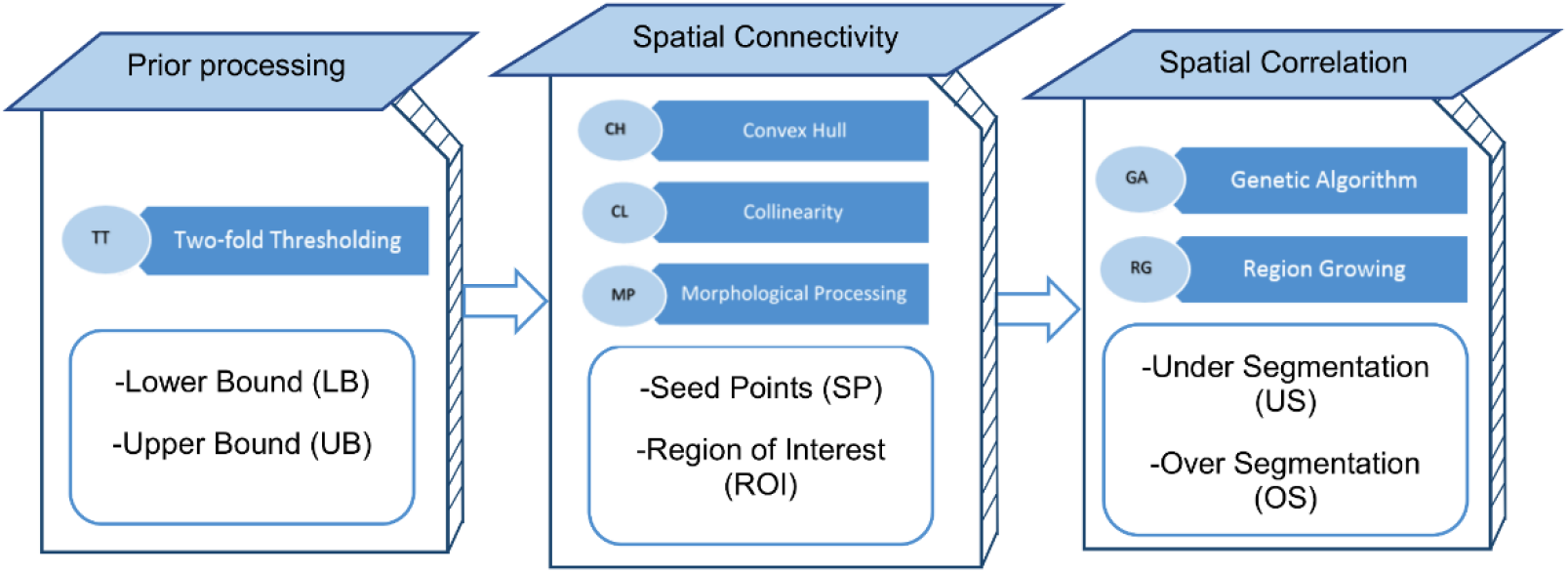
A three-layer architecture of spatial based image segmentation

Subsequently, a geometrical processing pipeline was applied to obtain a series of initial positions. Convex hull polygons were constructed on M_LB_ images and later masked onto the M_UB_ volume to produce a list of potential ROIs. The generated ROIs required refinement after the convexity and masking processes. Small-sized regions were treated as noise and discarded using the connected component labeling method. Only ROIs size larger than 200 pixels left intact. Afterward, a list of candidate seed points were created on these ROIs using collinearity property as visualized in Fig. 3 and described in stepwise manner below:

Step 1: Boundary tracing was applied to the polygons, and a list of vertices (*Xn* and *Yn*) was generated.
Step 2: The vertices were sorted based on the *Y* values.
Step 3: The vertices were sorted based on the *X* values.
Step 4: The middle point (mdp) was identified.

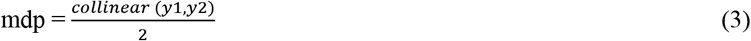
Step 5: The deepest point or convex hull point (chp) was located.

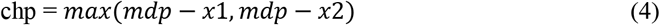
Step 6: Seed point 1 was located between mdp and chp.

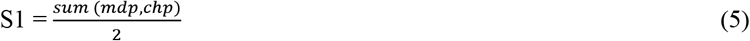
Step 7: Seed point 2 was located between chp and S1.

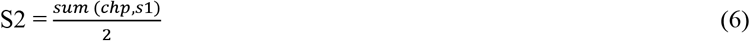
Step 8: Lastly, seed point 3 was located between mdp and S2.

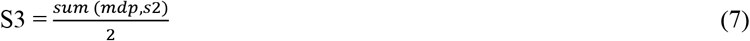

**Fig. 3.**
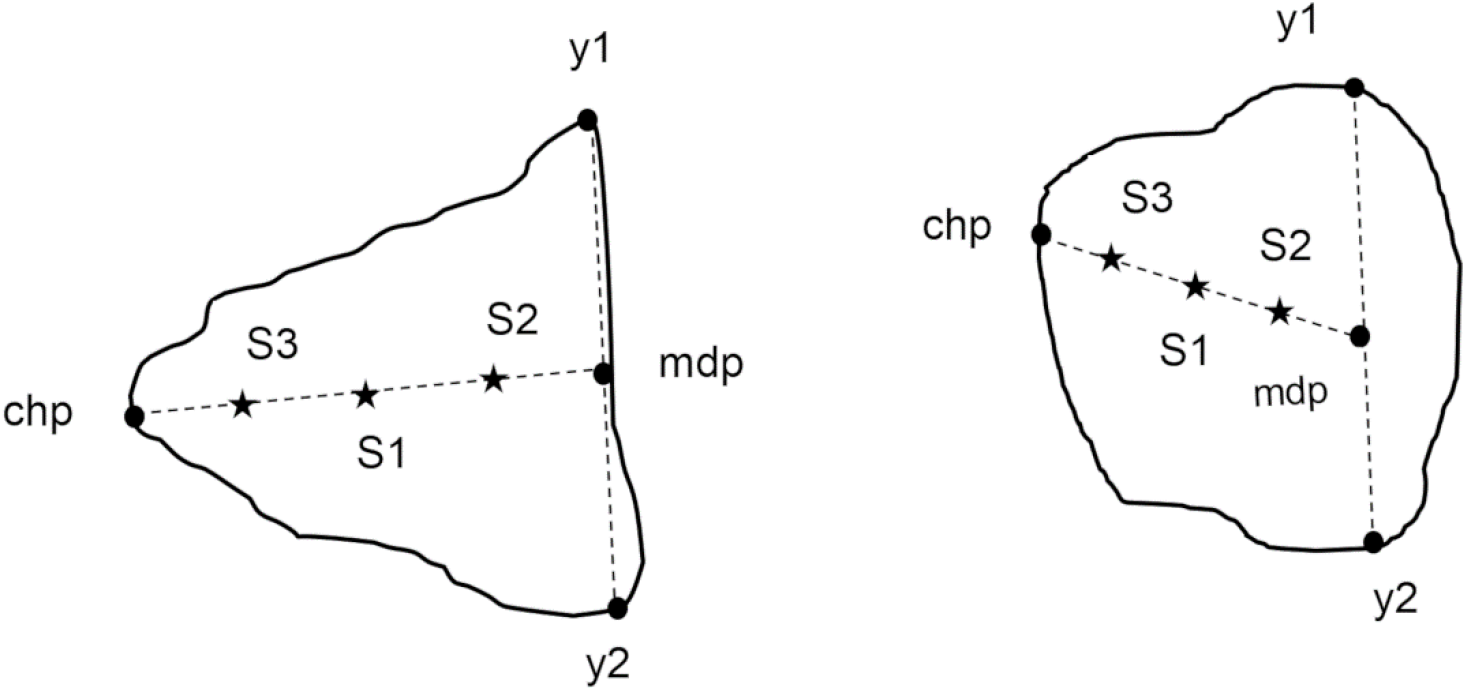
Example of seed points, S1, S2, and S3, were generated for each region of interest.

Finally a topological processing based on evolutionary region growing (ERG) [14–16] was applied to recognize the spatial relationship, which in this case was the homogeneity between groups of pixels. ERG is a robust method that can successfully rectify a distinct region growing issue such as leakage caused by sensitivity to the threshold, locality due to different initial points providing different results, and sensitivity to noise. A 15-bit binary chromosomes is designed. The first nine bits (*i = 9*) were assigned to the seed point candidates determined according to section 2.2.2; *S* = {*S_1_*, *S*_2_…*S_i_*}. The last six bits (*j = 15*) were assigned to the maximum distance variations allowed during region growth (T_7_ = 10, T_8_ = 20, T_9_ = 30, T_10_ = 40, T_11_ = 50, and T_12_ = 60). The following variables, settings, and initializations were fixed during region growth in ERG:

> Population size = 30, generation = 60, crossover operator = scattered, selection operator = roulette, mutation operator = uniform, crossover fraction = 0.8, mutation fraction = 0.2, and elitism = 0.1* population size.

### 2.3 Performance evaluation

A spatial overlapping metrics was constructed to measure the proposed framework performance as shown in Fig.4. Sensitivity represents the overlap between the ground truth and algorithm-detected regions. Specificity measures the non-overlapping pixels correctly excluded by the proposed algorithm. Accuracy considers both the overlapping and non-overlapping estimations. All three metrics were defined using hierarchical fitness functions as derived by Eq. 8, 9, and 10, respectively.

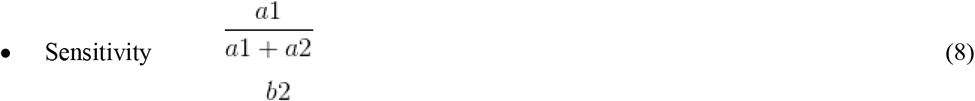

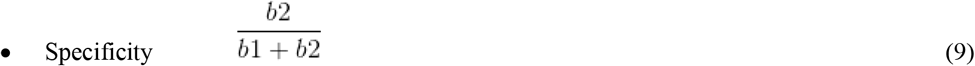

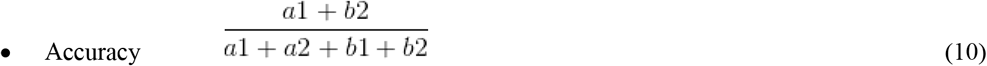

**Fig. 4.**
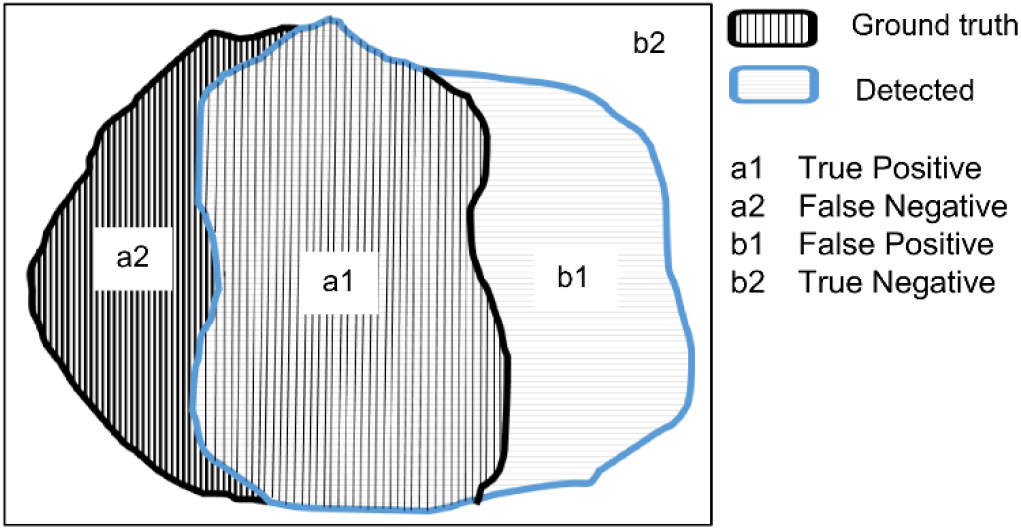
A true positive is defined as the overlap of pixels between the ground truth and region detected using the proposed method, whereas a false negative as the proposed method that fails to include a ground truth region. A true negative is defined as the exclusion of a region by the ground truth and proposed method, and a false positive is defined as a region that has been incorrectly included by the proposed method.

## 3. Results

The proposed algorithm converged and produced an optimal chromosome comprising the threshold and seed point information, which led to optimal segmentation. Table 2 presents the values of sensitivity, specificity, and accuracy, which evaluate the agreement of spatial overlap between the pixels and the quality of the segmentation. Sections 3.1–3.5 demonstrate examples of the corresponding slice-by-slice segmentation using the proposed method versus the ground truth, i.e., manual delineation by an expert.

**Table 2.**
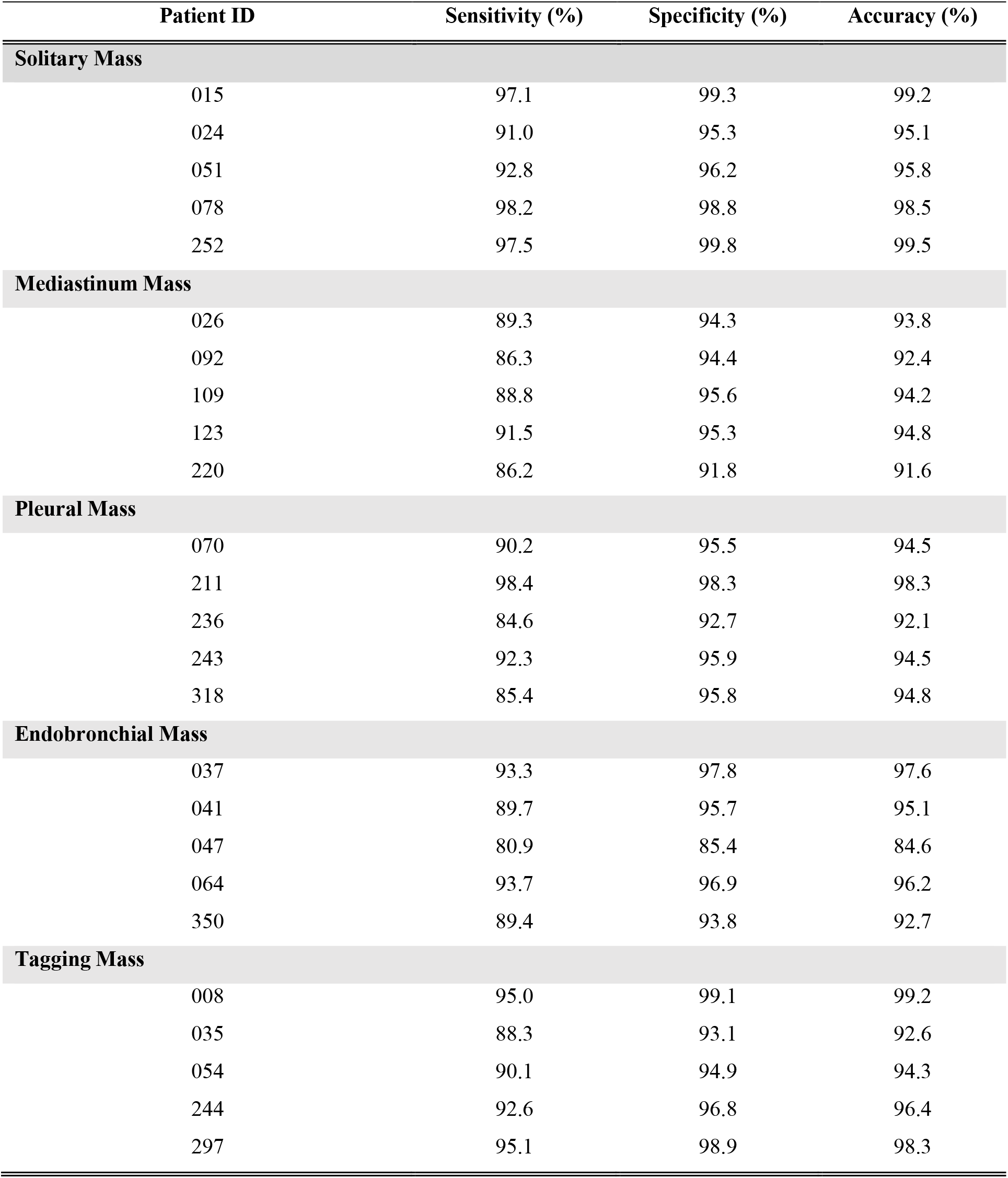
Performance metrics for each patient. Visualizations are provided in section 3.1–3.5 for all patients.

### 3.1 Solitary mass segmentation

Solitary masses independently exist (i.e., without other pathological objects) inside the lung parenchyma without invading surrounding structures. Accordingly, the assumption that these objects would require a less complicated segmentation was proven by the performances listed in Table 1, with sensitivity values of 91%–99% and specificity and accuracy values of 95%–99% each. Fig. 5 presents an example of solitary mass segmentation.

**Fig. 5.**
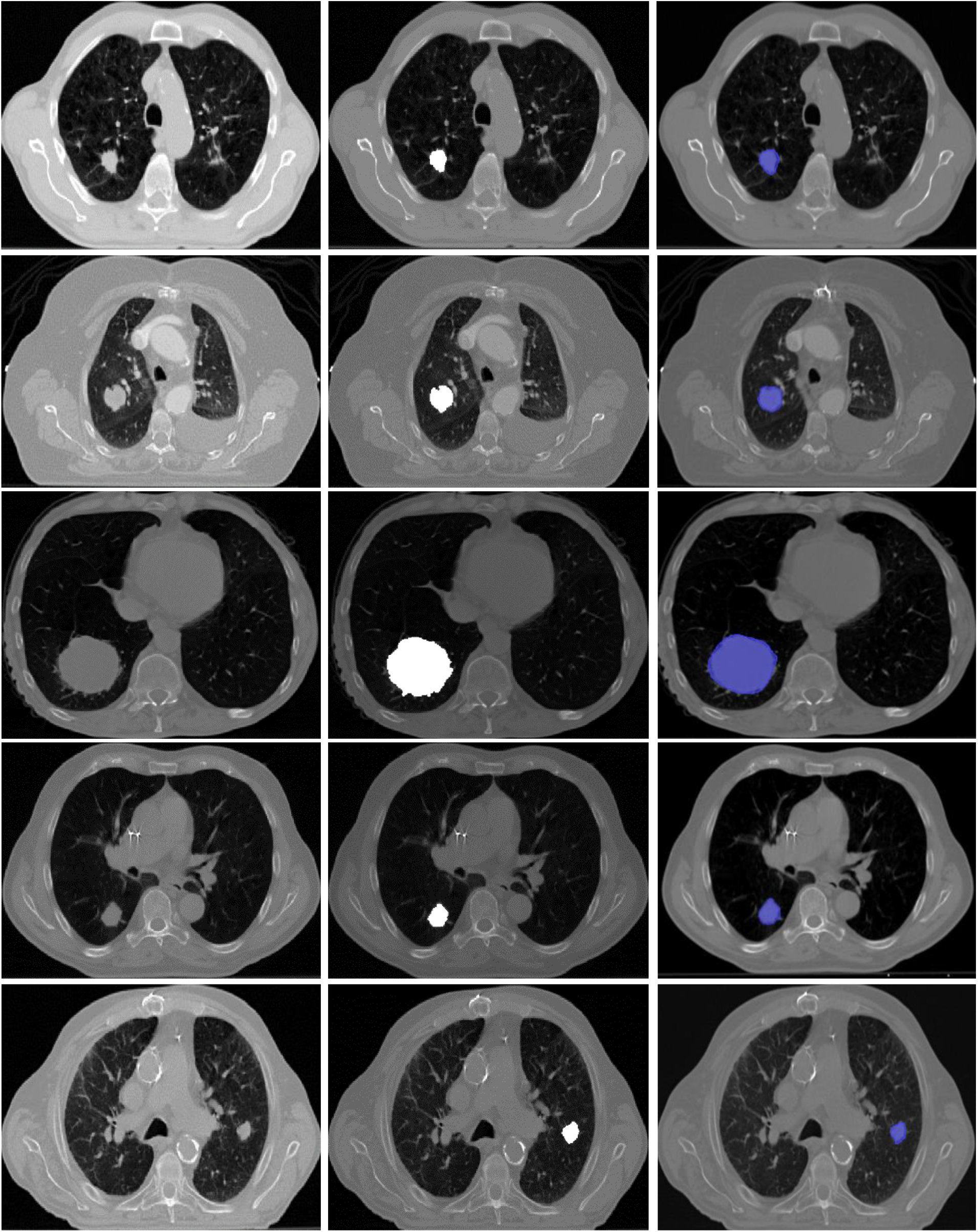
Solitary masses segmentation. The rows represent different patient ID (015, 024, 051, 078 and 252) from top to bottom. In contrast the columns represent raw images, proposed method results, and ground truth images, from left to right.

### 3.2 Mediastinal mass segmentation

Mediastinal masses often attach to the central compartment of the thoracic cavity between two pleural sacs (lobes) and may be connected to complicated structures such as the esophagus, trachea, thymus, and/or aorta. Therefore, mediastinal mass segmentation was a challenging task in this study, with sensitivity, specificity, and accuracy values of 86%–91%, 94%–96%, and 92%–96%, respectively. Fig. 6 presents an example of mediastinal mass segmentation.

**Fig. 6.**
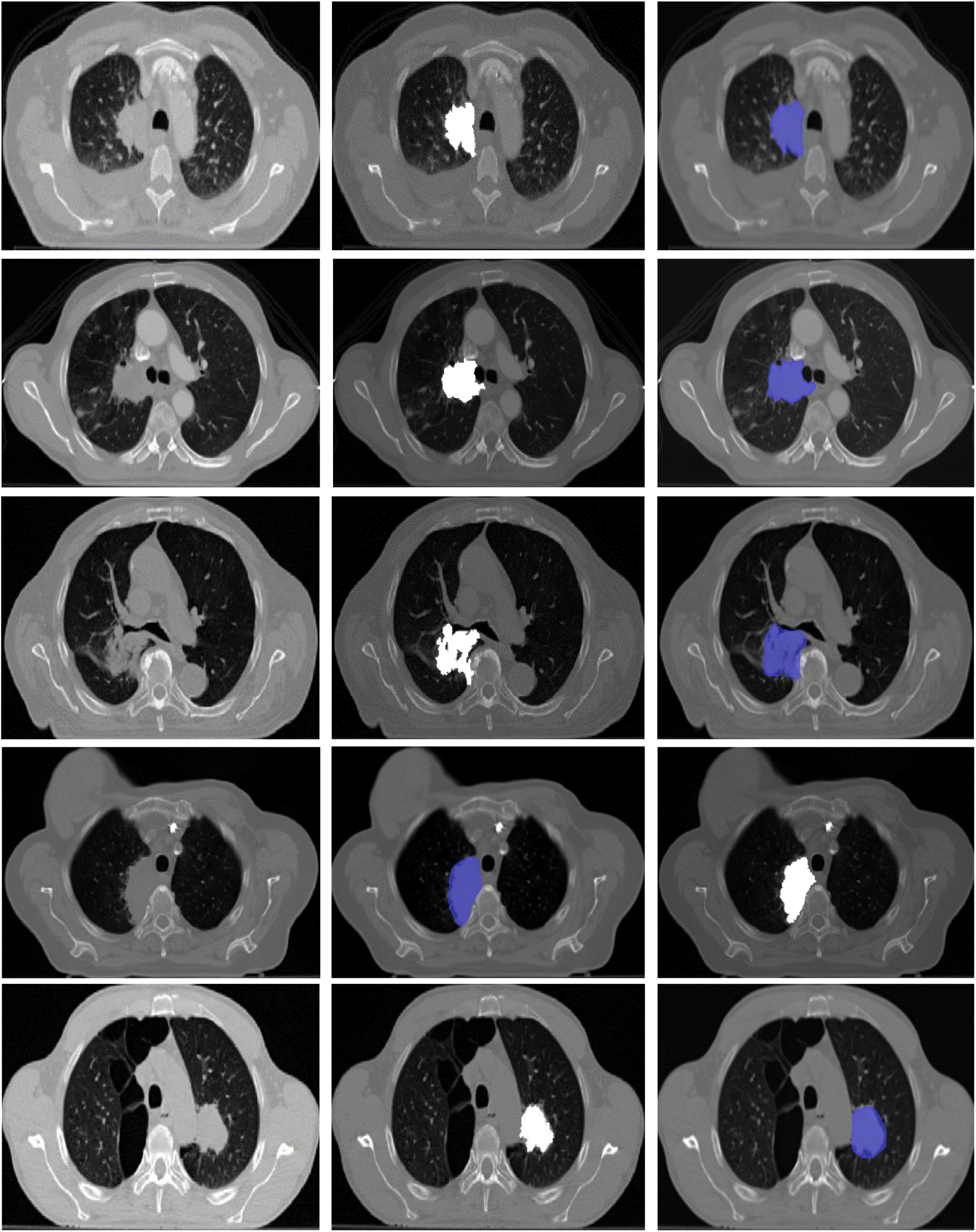
Mediastinum masses segmentation. The rows represent different patient ID (026, 092, 109, 123 and 220) from top to bottom. In contrast the columns represent raw images, proposed method results, and ground truth images, from left to right.

### 3.3 Pleural mass segmentation

Pleural masses invade the ribs or chest wall. Segmentation of these lesions was found to be less complicated than that of mediastinal masses because the former rarely exhibited tumor-like conditions (e.g., pathological objects that may reduce accuracy) but were more complicated than that of solitary masses. For pleural masses, sensitivity values of 85%–98% and specificity and accuracy values of 92%–98% each were obtained. An example of pleural mass segmentation is shown in Fig. 7.

**Fig.7.**
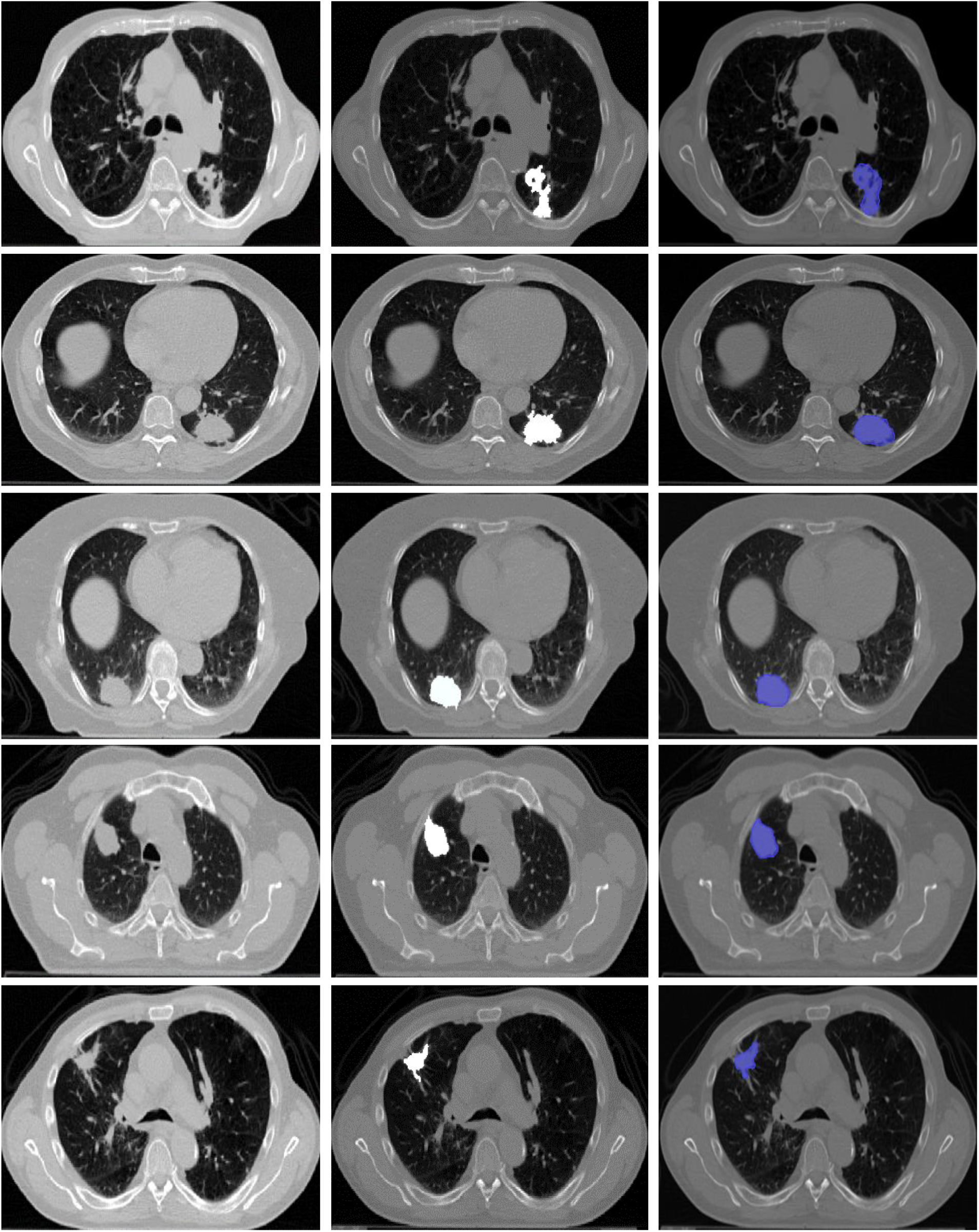
Pleural masses segmentation. The rows represent different patient ID (070, 211, 236, 243 and 318) from top to bottom. In contrast the columns represent raw images, proposed method results, and ground truth images, from left to right.

### 3.4 Endobronchial mass segmentation

Tracheo-bronchial tree lesions generally localize around the respiratory tract. These uncommon lesions are commonly classified as malignant. Like mediastinal masses, the segmentation of endobronchial masses was challenging, particularly because it was difficult to differentiate these lesions from the surrounding airway structures that exhibited similar intensities. Here, the sensitivity, specificity, and accuracy values was within the range of 80%– 93%, 85%–97%, and 84%–97%, respectively. Fig. 8 presents an example of mediastinal mass segmentation.

**Fig. 8.**
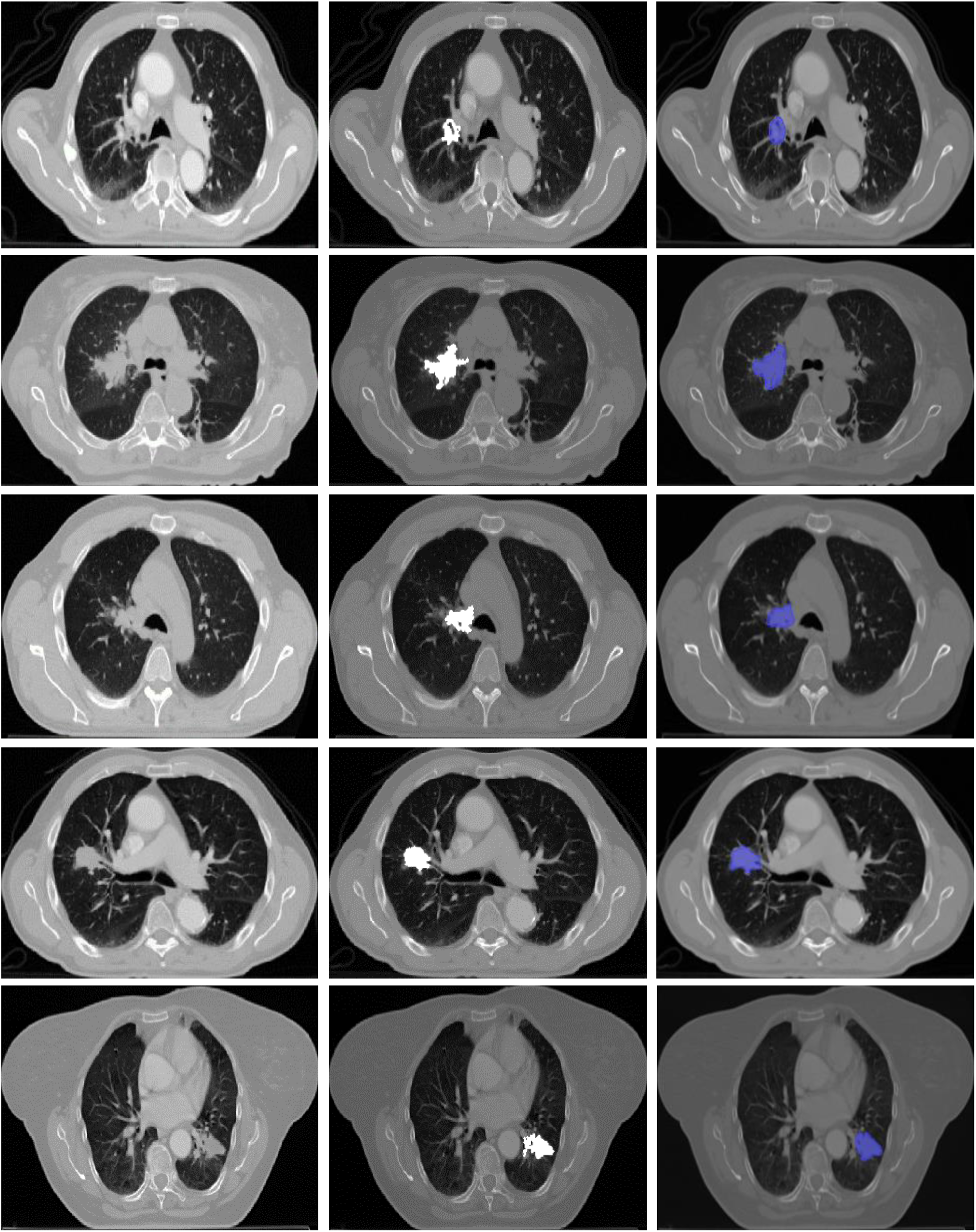
Endobronchial masses segmentation. The rows represent different patient ID (037, 041, 047, 064 and 350) from top to bottom. In contrast the columns represent raw images, proposed method results, and ground truth images, from left to right.

### 3.4 Tagging mass segmentation

Tagging masses are lesions that characteristically attach themselves to >1 of the following: pleura, mediastinum, or tracheobronchial tree. For these masses, the observed sensitivity, specificity, and accuracy values were 88%–95%, 93%–99%, and 92%–99%, respectively. An example of this type of mass segmentation is demonstrated in Fig. 9.

**Fig. 9.**
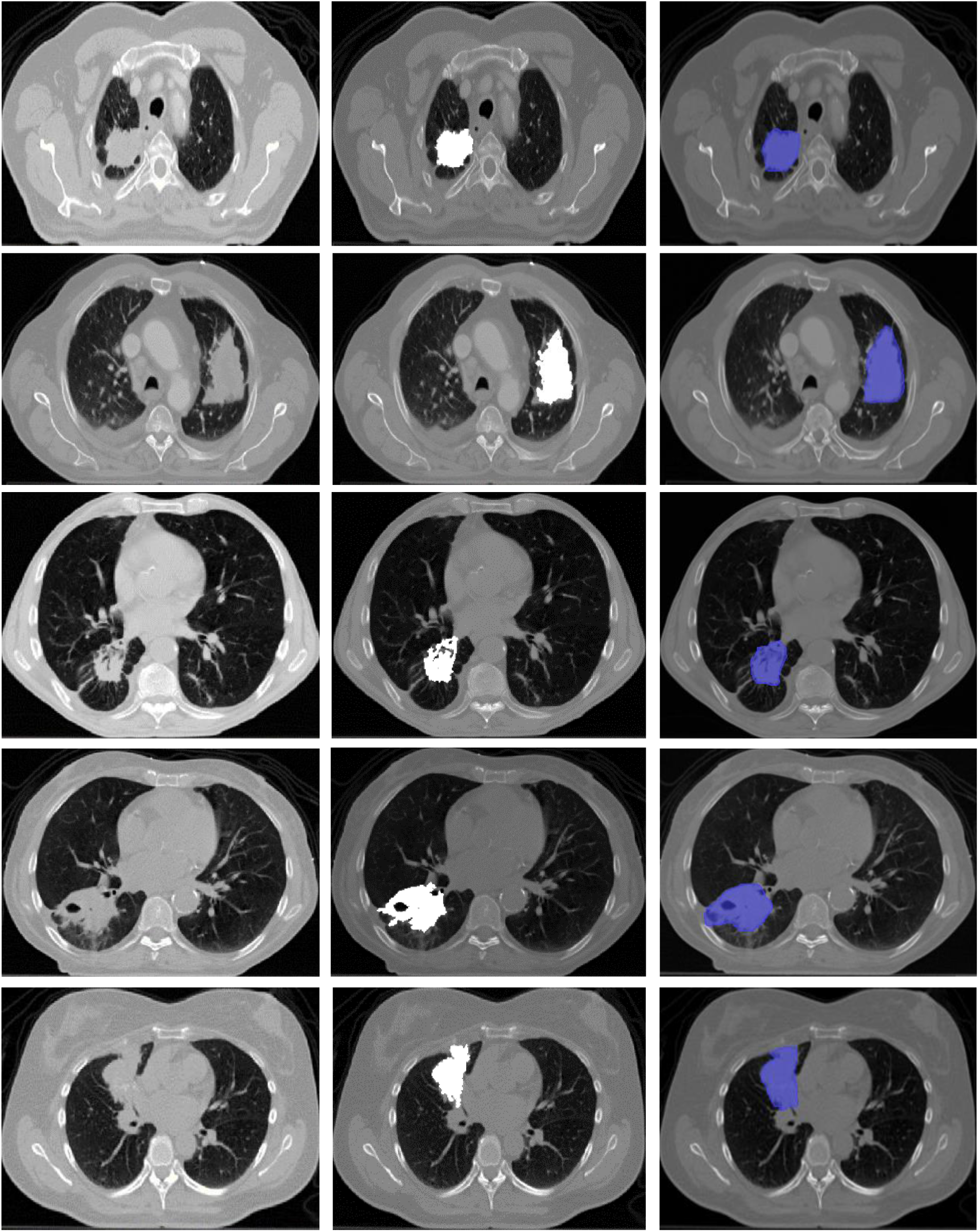
Tagging masses segmentation. The rows represent different patient ID (008, 035, 054, 224 and 297) from top to bottom. In contrast the columns represent raw images, proposed method results, and ground truth images, from left to right.

## Discussion

By invading the surrounding structures, such as blood vessels, chest and mediastinal walls, the NSCLCs may occlude the existent boundaries of the lungs, which increases the difficulty of accurately segmenting the lung thoracic areas on the chest CT images and hinders the subsequent detection and segmentation of lung cancers [17]. The existing algorithms for delineating lung nodules are also insufficiently efficient to segment the relatively larger NSCLC masses [3–7]. To rectify this problem, an algorithm was proposed that acknowledges the radio-morphological condition of a NSCLC. In this report, the robustness of the proposed algorithm was demonstrated in terms of the tumor size, shape (spherical or non-spherical), microenvironment (left or right lobe), localization (solitary, pleural, mediastinal, endobronchial, or tagging), and contouring (lobulated, cavitated, or spiculated) and have used 25 sets of medical (CT) images with these radio-morphological characteristics to evaluate the capability of the proposed algorithm.

For a minimum of four types of masses (solitary, pleural, mediastinal, and tagging), little to no difference was observed in comparison with the algorithm and. ground truth delineations. The lowest sensitivity values were expectedly obtained for endo-bronchial masses, which are considered the most complicated and difficult to extract even by expert radiologists. In such cases, the algorithm had to segment only the nodule while omitting the bronchus and other unrelated pathological objects attached to the same branch.

Regarding location, the algorithm easily extracted the ROI, regardless of the affected lobe side. Additionally, under-segmentation was observed only in one experimental case involving a mediastinal mass (Appendix A), which was assumed to be a unique event involving the “bridging” condition as shown. Here, bridging refers to a tumor with multiple connected regions that are separated by a shallow gap (i.e., change in intensity level), and this event ceased the expansion of the algorithm and overlooked the other region. Future research might address this condition by registering the tumor as having two regions. This bi-regional analysis would require a different seed-finding approach that initially treated the tumor as multiple separate regions, followed by co-registration at a later point.

Currently, the medical community relies on manual or semi-automatic tumor delineation for cancer diagnosis and detection, which is time-consuming and subject to fatigue and interobserver variability. In this study, a completely automated approach is presented intended to ease the burden of manual delineation while providing physicians with several options to select the most accurate delineation by determining a few seeds within the same suspicious ROI. Despite its advantages, however, the proposed method seems to be unable to avoid issues of explosion or over-segmentation, particularly in larger tumors. For such tumors, limiting the threshold value may cause under-segmentation, whereas increasing this value may cause explosion. The algorithm also failed to recognize a very large tumor that had expanded to both sides of the lung wall (Appendix B).

However, to the best of my knowledge, this is the first study to demonstrate a spatial-based segmentation algorithm for tumor detection and extraction. Several previous studies similarly discussed the use of evolutionary-based region growth to target lung parenchyma or thoracic region segmentation [14–16], but largely failed to address the tumor itself, which is the most vital part of medical image analysis. In contrast, the integration of spatial connectivity information with the backbone of evolutionary-based region growing algorithm in facilitating the tumor extraction seems promising. The favorable results obtained in diverse radiological situations demonstrate the potential of the proposed method.

## Conclusion

With this study, the requirement of a robust, completely automated tumor extraction method intended to reduce the burdens on clinicians during medical image analysis have been addressed. To that end, a potential spatial information-based segmentation method that hybridizes spatial connectivity and spatial relationships between pixels to detect and extract ROIs was demonstrated. The proposed method will provide beneficial assistance in the clinical diagnosis, staging, and therapy response assessment of cancer patients.

## Acknowledgments

I am grateful to Dr. Norafida Bahari (Radiologist and Senior Medical Lecturer in the Department of Imaging, Faculty of Medicine and Health Sciences, University Putra Malaysia) for her assistance on the ground truth work.

## Ethical approval

The clinical materials were taken from a publicly available database, The Cancer Imaging Archive (TCIA), which were made available for download by the Dana–Farber Cancer Institute in accordance with the Dutch law and Institutional Review Board approval. The Institutional Review Board of the Maastricht University Medical Center (MUMC+) waved review due to the retrospective nature of the original study.

## Appendix A

**Fig. A.**
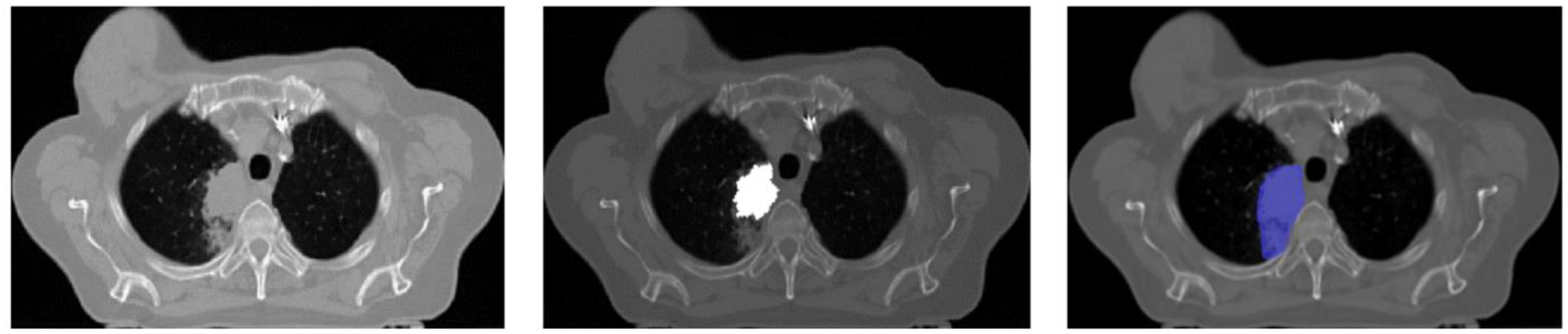
Under segmentation case on patient 123 which happened on one of the slices.

## Appendix B

**Fig. B.**
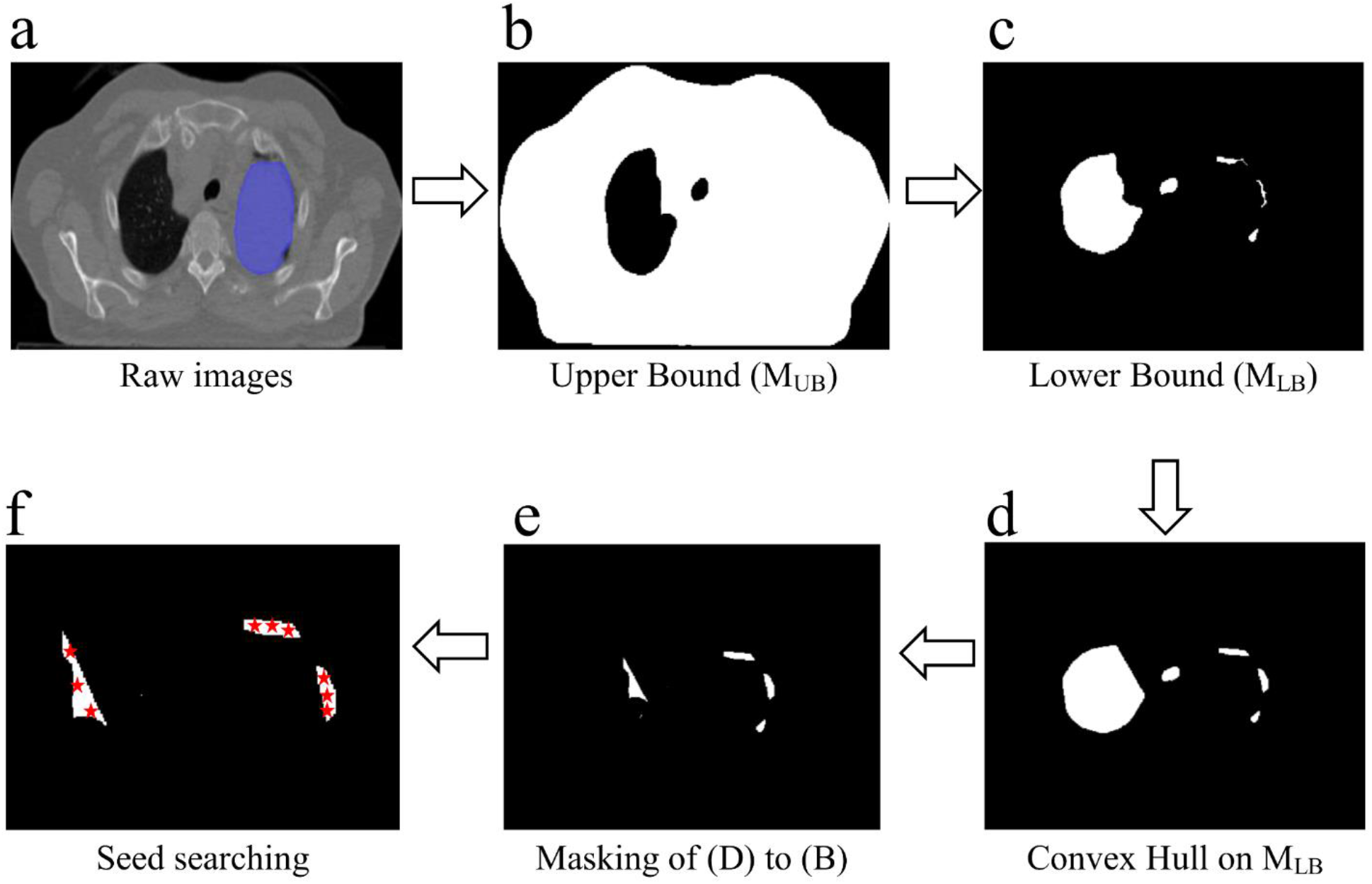
An example of a case where the proposed algorithm failed to work. The tumor almost entirely covers the left lobe; therefore, during the two-fold thresholding as shown in (B), the target region could not be detected. Note that the candidate ROIs in (E), where seeds to be generated excluded the real tumor region, hence the algorithm failed in such event.

## References

[1] Seow P, Win MT, Wong JHD, Abdullah NA, Ramly N (2016) Segmentation of solid sub region of high grade gliomas in MRI images based on active contour model (ACM). J Phys: Conf Ser 694:012043. doi:10.1088/1742-6596/694/1/012043

[2] Hu YC, Grossberg MD, Wu A, Riaz N, Perez C, Mageras GS (2012) Interactive semiautomatic contour delineation using statistical condition random fields framework. Med Phys. 39(7):4547–4558. doi: 10.1118/1.4728979.

[3] Farag AA, HE EL Munim, Graham JH, Farag AA (2013) A novel approach for lung nodules segmentation in chest CT using level sets. IEEE Transactions on Image Processing. 22(12):5202–5213.

[4] Messay T, Hardie RC, Tuinstra TR (2015) Segmentation of pulmonary nodules in computed tomography using a regression neural network approach and its application to the Lung Image Database Consortium and Image Database Resource Initiative dataset. Med Image Anal. 22(1):48–62. doi.org/10.1016/j.media.2015.02.002

[5] Elmar RZ, Ponomaryov V (2016) Automatic lung nodule segmentation and classification in CT images based on SVM. In Physics and Engineering of Microwaves, Millimeter and Submillimeter Waves (MSMW). 9th International Kharkiv Symposium on. 1–4.

[6] Hu Y, Menon PG (2016) A neural network approach to lung nodule segmentation. Medical Imaging: Image Processing. pp 97842O. doi: 10.1117/12.2217291

[7] Krishnamurthy S, Narasimhan G, Rengasamy U (2016) Three-dimensional lung nodule segmentation and shape variance analysis to detect lung cancer with reduced false positives Proc Inst Mech Eng H. 230(1):58–70. doi: 10.1177/0954411915619951

[8] Whitfield GA, Price P, Price GJ, Moore CJ (2013) Automated delineation of radiotherapy volumes: are we going in the right direction. Br J Radiol. 89(1021):20110718. doi: 10.1259/bjr.20110718

[9] Wu K, Ung YC, Hwang D, Tsao MS, Darling G, Maziak DE, Tirona R, Mah K, Wong CS (2010) Autocontouring and manual contouring: Which is the better method for target delineation using F-DG PET/CT in Non-Small Cell Lung Cancer? J Nucl Med. 51(10):1517–23. doi: 10.2967/jnumed.110.077974

[10] Korsager AS, Carl J, Ostergaard LR (2016) Comparison of manual and automatic MR-CT registration for radiotheraphy of prostate cancer. J Appl Clin Med Phys. 7(3):294–303. doi: 10.1120/jacmp.v17i3.6088

[11] Clark K, Vendt B, Smith K, Freymann J, Kirby J, Koppel P, Moore S, Phillips S, Maffitt D, Pringle M, Tarbox L, Prior F (2013) The cancer imaging archive (TCIA): Maintaining and operating a public information repository. J. Digital Imag. 26(6):1045–1057. doi: 10.1007/s10278-013-9622-7

[12] Aerts HJWL, Velazquez ER, Leijenaar RTH, Parmar C, Grossmann P, Carvalho S, Lambin P (2015) The Cancer Imaging Archive. http://doi.org/10.7937/K9/TCIA.2015.PF0M9REI

[13] Mesanovic N, Grgic M, Huseinagic H, Males M, Skejic E, Smajlovic M (2011) Automatic CT image segmentation of the lungs with region growing algorithm. 18th International Conference on Systems, Signals and Image Processing—IWSSIP. 395–400.

[14] Allaoui AEL, Nasri M (2012) Threshold optimization by genetic algorithm for segmentation of medical images by region growing. International Journal of Emerging Trends and Technology in Computer Science. 1(2):161–166.

[15] Allaoui AEL, Merzougui M, Mirhisse J (2016) Evolutionary algorithm for segmentation of medical images by region growing Computer Graphics. Imaging and Visualization (CGiV). 13th International Conference on 2016 Mar 29. 119–124. doi: 10.1109/CGiV.2016.32

[16] Gowri BS, Ilango G (2014) Segmentation of medical images using topological concepts based region growing method. J Math. 10(4):1–7

[17] Zhou J, Chang S, Liu Q, Metaxas DN, Zhao B, Ginsberg MS, Shwartz LH (2008) Automatic detection and segmentation of large lung cancers from chest CT images. Workshop of 11th international conference on medical image computing and computer assisted intervention. 165–174.

